# Quorum sensing employs a dual regulatory mechanism to repress T3SS gene expression

**DOI:** 10.1101/2024.07.30.605901

**Authors:** Payel Paul, Ram Podicheti, Logan J. Geyman, Kai Papenfort, Julia C. van Kessel

## Abstract

The type III secretion system (T3SS) is a needle-like complex used by numerous bacterial pathogens in host infection by directly injecting exotoxins into the host cell cytoplasm, leading to cell death. The T3SS is a known virulence factor in the shrimp pathogen *Vibrio campbellii*. The ∼40 genes comprising the *V. campbellii* T3SS are regulated by a network of transcription factors in response to changes in the cell’s environment: cell density (quorum sensing; QS), temperature, calcium, and host cell contact. Under positive environmental stimuli, the master T3SS transcription factor ExsA activates expression of the four structural T3SS operons required for needle formation. Previous studies identified a key role of the master QS transcription factor LuxR: repression of *exsA* transcription via DNA binding at the *exsBA* promoter. Here we uncovered a new regulatory role of LuxR: indirect post-translational repression of ExsA activity via direct transcriptional repression of the gene encoding the anti-anti-activator ExsC. In *V. campbellii*, ExsC is a positive regulator of T3SS transcription: high ExsC expression leads to full ExsA transcription activation of the T3SS structural promoters. LuxR binding at the *exsC* promoter represses transcription of *exsC* through disruption of ExsA binding. Our findings collectively show that *V. campbellii* responds to high cell density signals to shut down the expression of the T3SS. We postulate that this dual regulatory mechanism by LuxR enables both the rapid inactivation of existing ExsA protein and blocks its further synthesis, leading to a rapid shutdown of T3SS activity at high cell density.

**Importance:** *Vibrio campbellii* utilizes the type III secretion system (T3SS) as a mechanism of pathogenesis, which is a highly studied ‘injectisome’ complex that delivers exotoxins into host cells during infection. The T3SS pathogenicity island in *V. campbellii* comprises ∼40 genes that are organized into four structural operons. In this study, we determined that quorum sensing – a method of bacterial communication – regulates T3SS genes both at the transcriptional and post-translational levels to shut down T3SS gene expression at high population densities.

## INTRODUCTION

*Vibrio campbellii* is a Gram-negative γ-proteobacterium that belongs to the class *Vibrionaceae*, many of which are recognized as pathogens of aquatic animals, including shrimp, lobster, mollusk, shellfish, *etc.* (1–5). Pathogenic strains of *V. campbellii* are known to cause diseases in shrimp such as acute hepatopancreatic necrosis disease (AHPND), red spot syndrome, white tail disease, and luminous vibriosis among others (1, 3, 6–8). In our study, we use *V. campbellii* BB120 (*a.k.a.*, ATCC BAA-1116, previously classified as *V. harveyi*) (9) as a model organism to study mechanisms of virulence in pathogenic strains of *V. campbellii*.

A major mechanism of virulence in Gram-negative pathogens is the type III secretion system (T3SS) machinery, a specialized secretion apparatus which plays an essential role in secretion of extracellular protein toxins. It is a syringe-like membrane-embedded apparatus also called the ‘injectosome’, which contacts the eukaryotic host cell membrane and injects effector proteins or exotoxins across the membrane and into the host cell (10, 11). Exotoxins compromise the host cell machinery and activate cell lysis through various mechanisms that have been broadly studied in pathogens including *Yersinia pestis*, *Pseudomonas aeruginosa*, *Salmonella typhimurium*, and *Vibrio parahaemolyticus* (12–16). Certain species of the *Vibrionaceae* family such as environmental isolates of *V. parahaemolyticus* possesses two types of T3SSs encoded by entirely separate sets of structural genes and carrying distinct arsenals of effector proteins (11). The two T3SSs are broadly differentiated based on the target and general mechanism of action of the effector proteins. While T3SS1 encodes effector proteins responsible for cytotoxicity in host cells and survival of bacterial cells in the host environment, the T3SS2 encodes for protein toxins responsible for host cell invasion and seafood borne gastroenteritis in humans (15). T3SS2 is also found in some environmental strains of *V. cholerae* (11).

Previous studies identified a functional T3SS in *V. campbellii* BB120 and determined that QS negatively regulates T3SS at high population densities (17–20). Further, QS-mediated regulation of T3SS has been shown to be an important virulence factor in *V. campbellii* infection in shrimp (21). All data thus far indicate that the T3SS of *V. campbellii* BB120 is similar in structure, function, and genetic organization to the T3SS1 of *V. parahaemolyticus*; *V. campbellii* BB120 does not encode a T3SS2 (18, 20). Thus, we focus here on T3SS1 function and regulation in *V. campbellii*.

Bacterial pathogens regulate the expression of their virulence traits – such as T3SS – in response to external signals to initiate colonization of target tissues, avoid host immune system mechanisms, and persist. In *V. parahaemolyticus and P. aeruginosa,* the Exs proteins ExsA, ExsD, ExsC, and ExsE establish a regulatory cascade that dictates the activity of the T3SS1 master transcription activator ExsA (22–28) (Fig. 1A). In non-inducing conditions, *i.e*., the absence of a host cell membrane, or in media containing high levels of calcium and minimal magnesium, ExsA is bound by its anti-activator ExsD, and the T3SS genes are not transcribed by ExsA. Also in this condition, chaperone ExsC binds secretory protein ExsE. ExsE is not secreted while T3SS channels remain closed. Upon sensing inducing conditions, *i.e*., upon contact with a host cell membrane or the absence/decrease of calcium and addition of magnesium, T3 secretion is induced, allowing ExsE to be secreted by the T3SS (Fig. 1A). Once ExsE is secreted, ExsC is free to bind its other binding partner ExsD, which relieves the repression of ExsA. ExsA is then free to bind the downstream T3SS1 promoters and activate their transcription (26, 27, 29). Induction of T3SS genes greatly increases the rate of synthesis of T3SS machinery. In addition to these four factors, the *exsB* gene clusters with the regulatory *exs* genes. It is a pilotin that attaches to the outer membrane and regulates the assembly of the type III secretin on the outer membrane (30). Collectively, the Exs regulatory cascade composes the internal post-translational regulatory system.

**Figure 1.**
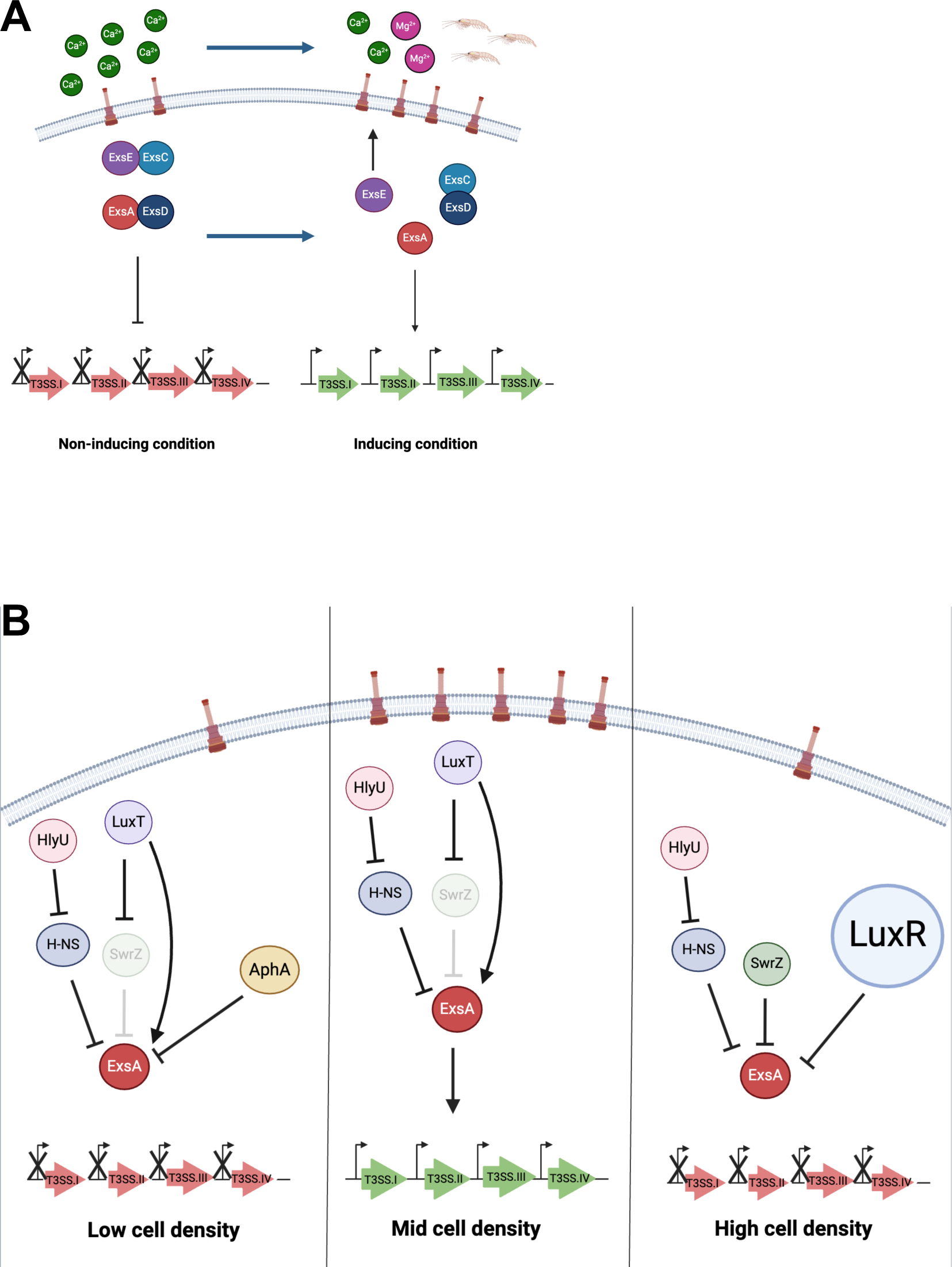
Schematic models of the previously determined regulatory networks that control T3SS gene expression. Both diagrams show the current model in the T3SS field in which secretion is induced on contact with host cell membrane during infection, or on chelation of calcium ions from media along with addition of magnesium. **(A)** The schematic shows the internal Exs regulatory cascade determined in *Pseudomonas*, which comprises ExsA, ExsD, ExsC and ExsE. The effect of these proteins on the expression of T3SS structural operons I - IV is indicated. **(B)** The schematic shows the defined regulators of T3SS genes that have been determined in *V. campbellii* BB120, which include quorum sensing regulators LuxR and AphA, H-NS, LuxT, and SwrZ. LuxT is a direct and indirect activator of *exsA* at low cell density (LCD), AphA is a repressor of *exsA* at LCD, and LuxR is a repressor of *exsA* at high cell density (HCD). The roles of HlyU, H-NS, and SwrZ at different cell densities have not yet been determined, thus these are included in all three density stages. T3SS structural genes are turned on at mid cell density (MCD) and are off at LCD and HCD.

The *Vibrio* T3SS1 operons are also regulated at the transcriptional level by a wide variety of transcription factors and nucleoid binding proteins. These include: 1) H-NS, a histone-like nucleotide binding protein and global regulator of gene expression that represses T3SS transcription by binding along the length of the promoter regions of T3SS1 genes *exsE*, *exsC*, *exsB*, and *exsD* (26, 31); 2) HlyU, a global activator of virulence genes that relieves repression of DNA transcription by removing H-NS thereby acting as an activator of T3SS (32); 3) LuxT, a global low cell density (LCD) regulator which activates ExsA expression indirectly by repressing *swrZ* expression (17); 4) SwrZ, a regulator of swarming motility which represses ExsA expression (17); and 5,6) LuxR and AphA, QS regulators of high and low cell densities (HCD and LCD), respectively, which repress *exsA* transcription (19) (Fig. 1B). Thus, a local and global network of regulators act to limit T3SS expression and activation within a narrow window at mid cell density (MCD) (19).

Previous studies of QS regulation showed that LuxR inhibits T3SS gene expression at HCD through the direct binding of LuxR to the *exsB* and *exsA* promoters that drive expression of *exsA*, resulting in no production of the T3SS activator ExsA (19, 20). Further, LuxR primarily regulates *exsA* transcription through the *exsB* promoter; *exsA* and *exsB* are expressed as co-transcripts in *V. campbellii* (20). We observed that a strain in which *exsA* expression was decoupled from its native promoter retained a regulatory effect by LuxR on transcription of T3SS genes. Thus, we sought to test for other mechanisms of LuxR regulation of T3SS. In this study, we show evidence that LuxR binds and regulates transcription from both the *exsB* and *exsC* promoters in *V. campbellii* BB120, resulting in a dual mechanism of regulation: 1) LuxR represses *exsA* transcription via the *exsB* promoter (20), and 2) LuxR regulates ExsA activity post-translationally through repression of *exsC* transcription. We hypothesize that the dual regulation allows the QS system to shut-down the T3SS quickly and efficiently at high cell density.

## RESULTS

### A reporter assay for measuring T3SS regulation

To formally assess the predicted role of the T3SS regulatory genes in our hands, we designed a reporter system to enable rapid quantification of T3SS gene regulation. We constructed reporters with a promoterless *gfp* gene fused to one of two T3SS gene promoters, *exsD* or *vopN* (Fig. S1A, S1B). As expected from previous studies, LuxR repressed *exsD* and *vopN* expression, and ExsA was required for expression of both genes. The P*_vopN_* reporter yielded the larger dynamic range (Fig. S1A, S1B), and the fluorescence expression results were corroborated with RT-qPCR (Fig. S1C, S1D). Thus, we used this reporter construct moving forward. In addition, all assays measuring T3SS were induced in liquid medium with 15 mM MgSO_4_ to obtain more robust expression of T3SS genes as described in a previous study (33).

We first tested our *vopN* reporter in *V. campbellii* BB120 with genetically locked mutant strains that mimic the LCD and HCD conditions of QS. In *Vibrios*, the QS cascade involves multiple membrane-bound, histidine-kinase receptor proteins (Fig. S2) (34). In the absene of autoinducers at LCD, phosphorylation of the phospho-transfer protein LuxU by these receptor kinases (unbound by autoinducers) then phosphorylates the response regulator LuxO, which drives transcription of the Qrr regulatory RNAs and leads to LCD gene expression by AphA (35). Conversely, at HCD, autoinducer binding to the membrane-bound receptors reverses the phosphorylation cascade, resulting in unphosphorylated LuxO protein, no Qrr expression, and HCD gene expression by LuxR (19). Thus, we used genetically locked mutant strains to mimic the LCD and HCD states: *luxO* D61E and Δ*luxO* strains, respectively (36–40). Of note, it was discovered that previous annotations of *luxO* in *V. cholerae* and other *Vibrio* species were incorrect, and thus previous descriptions of the phosphomimic *luxO* D47E were actually *luxO* D61E (37). We also found this to be true for *V. campbellii* DS40M4, and thus refer to this allele as *luxO* D61E throughout this manuscript. The *luxO* D61E strain showed high levels of P*_vopN_-gfp* transcription, similar to the Δ*luxR* strain (Fig. 2A). Conversely, the Δ*luxO* strain showed diminished levels of P*_vopN_-gfp* transcription, similar to the wild-type strain at HCD (Fig. 2A). These patterns aligned with all previous studies of the QS regulatory epistasis (18–20).

**Figure 2.**
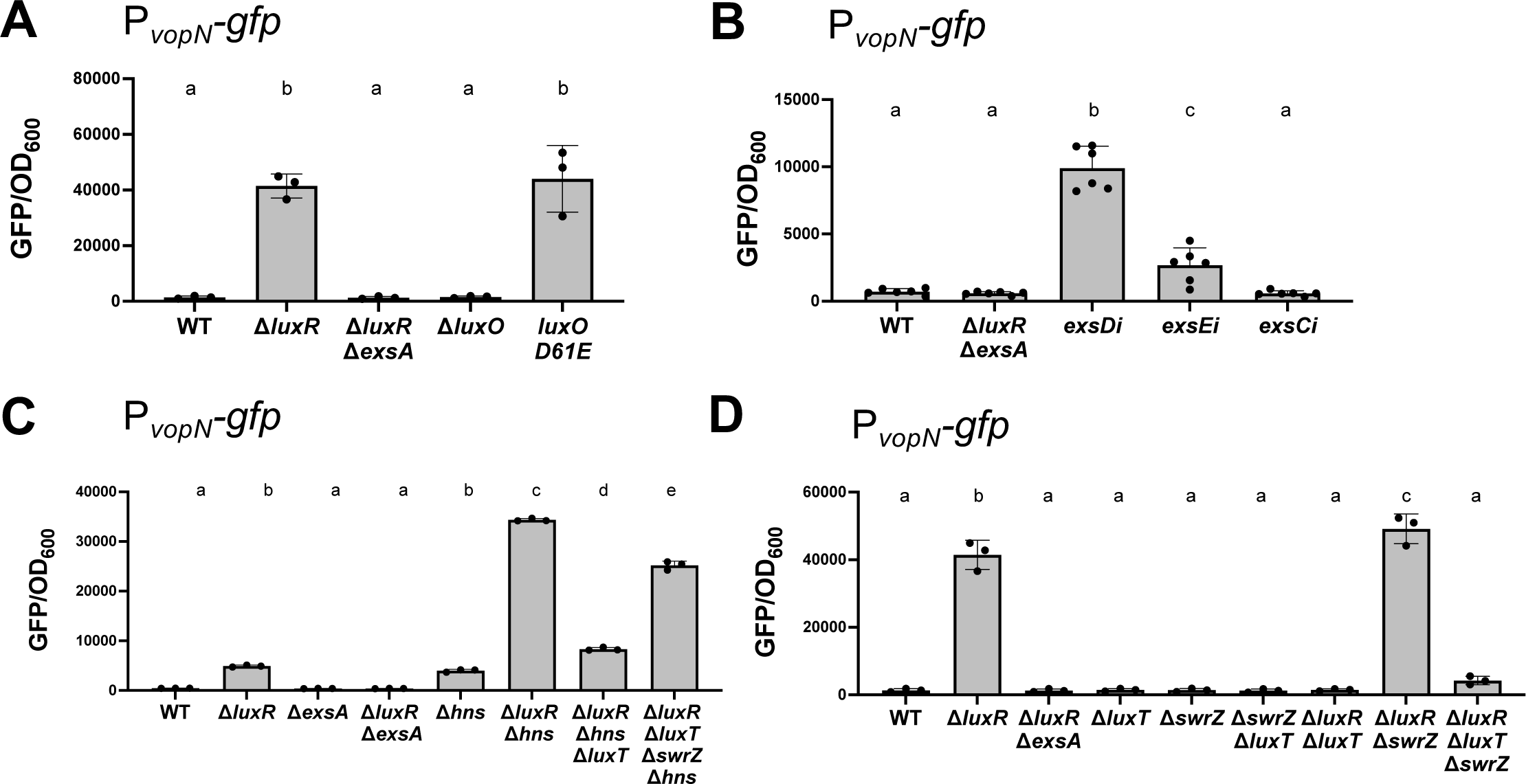
The P*_vopN_-gfp* reporter assay corroborates previous regulatory studies of T3SS genes in *V. campbellii.* **(A-D)** GFP reporter assays measure transcription from the *vopN* promoter in isogenic strains of *V. campbellii* BB120. **(B)** CRISPRi was used to knockdown expression of *exsD* (*exsDi*), *exsE* (*exsEi*), and *exsC* (*exsCi*) via induction of CRISPRi with 100 μM IPTG. For all panels, the error bars represent the standard deviation of means of *n*=3 biological replicates. Different letters indicate significant differences in pair-wise comparisons; the same letters indicate no significant differences (*p<*0.05). (one-way ANOVA followed by Tukey’s multiple comparisons test on normally distributed data (Shapiro-Wilk test), *p <* 0.01, *n* = 3).

Given the similarity of the *V. campbellii* T3SS to *V. parahaemolyticus*, we hypothesized that the Exs regulatory cascade comprising ExsA, ExsD, ExsC, and ExsE proteins regulate the T3SS in *V. campbellii* in the same manner as in *V. parahaemolyticus*. We tested homologs of the Exs proteins in *V. campbellii* for their regulatory roles by designing transcriptional knockdowns of the *exs* genes using CRISPRi technology (41). In the *exsD* and *exsE* knockdown strains, GFP fluorescence from the *vopN* promoter was significantly increased compared to wild-type (Fig. 2B). This result was expected because, if similar to the *V. parahaemolyticus* model, a decrease in ExsD protein would free ExsA protein to enable transcription activation, and a decrease in ExsE would free ExsC to interact with ExsD and thus release ExsA. In the *exsC* knockdown strain, GFP fluorescence and hence transcription from the *vopN* promoter was similar to wild-type (Fig. 2B). This result was expected because if the epistasis determined for *V. parahaemolyticus* is conserved in *V. campbellii*, a decrease in ExsC would result in more unbound ExsD to bind with ExsA and block transcription activation. We conclude that the effect of ExsC, ExsD, and knockdown in *V. campbellii* follows a pattern that is predicted by the *V. parahaemolyticus* system.

Next, we sought to examine the previously reported roles of global regulators H-NS, LuxT, and SwrZ in regulating the T3SS gene expression using our reporter assay. In *V. campbellii* and *V. parahaemolyticus*, H-NS has been shown to inhibit T3SS1-related gene expression through repression of *exsA* (26, 31). Our reporter assays corroborated these findings: GFP expression increased in the Δ*hns* mutant and was maximally expressed in the Δ*luxR* Δ*hns* strain (Fig. 2C). According to an earlier study, LuxT acts as an activator of T3SS genes at LCD and SwrZ represses T3SS gene expression (17). The authors concluded that LuxT activates T3SS through ExsA via de-repression of SwrZ as well as independently of SwrZ. Our assay confirmed that LuxT plays an important role in activating T3SS gene expression: deletion of *luxT* showed significantly decreased GFP expression in two strain backgrounds (Fig. 2C, 2D; compare Δ*luxR* Δ*hns* versus Δ*luxR* Δ*hns* Δ*luxT* or compare Δ*luxR* Δ*swrZ* versus Δ*luxR* Δ*swrZ* Δ*luxT*). In addition, deletion of *swrZ* in the Δ*luxR* background significantly increased GFP expression, although this was a modest increase (Fig. 2D). These results indicate that both LuxT and SwrZ indeed activate and repress T3SS expression, respectively, as previously shown. The large reduction in P*_vopN_* transcription in Δ*luxR* Δ*swrZ* Δ*luxT* compared to Δ*luxR* Δ*swrZ* was likely due to the SwrZ-independent activating role of LuxT described by Eickhoff *et al*. We wanted to test whether LuxT had an activating phenotype because it blocked H-NS or through another mechanism. We deleted *hns* in the Δ*luxR* Δ*luxT* Δ*swrZ* strain and observed that P*_vopN_* transcription significantly increased (Fig. 2C). However, transcription was not restored to the level of the Δ*luxR* Δ*hns* mutant, indicating that LuxT likely plays a role in activation of T3SS genes beyond the repression of SwrZ and H-NS activity (Fig. 2C). Collectively, these results show that the *vopN* reporter assay accurately reflects the genetic regulation of T3SS structural genes defined in previous studies and predicted by comparative genomics.

### LuxR regulates T3SS through the exsBA and exsC promoters

In *V. campbellii* BB120, the ∼40 genes composing the T3SS are on a pathogenicity island and are sub-divided into four structural operons called T3SS I, II, III, and IV with promoters in front of the *vcrG*, *vopN*, *vscN*, and *exsD* genes, respectively (Fig. S3A) (18). ExsA is required for activation of T3SS gene expression of all four structural operons (20). Given the role of LuxR as a negative regulator of T3SS activation, we wanted to test if LuxR regulates the T3SS exclusively through repression of *exsA* transcription from the *exsBA* operon or if there are additional T3SS loci regulated by LuxR. We hypothesized that LuxR represses transcription from the four T3SS structural operons in addition to shutting down *exsA* transcription from the *exsBA* operon at HCD (Fig. S3A). To test our hypothesis, we designed an assay in which *exsA* transcription and translation was uncoupled from its native promoter and T3SS gene expression was measured: 1) the strain background is Δ*exsA* Δ*exsB*, and *exsA* was expressed ectopically from a plasmid wherein transcription was under the control of an IPTG-inducible promoter (P*_tac_-exsA*), 2) translation of ExsA was also regulated through the placement of a theophylline-dependent riboswitch next to the promoter to regulate ribosome binding and translation (P*_tac-theo_-exsA*) (42, 43), and 3) the P*_vopN_-gfp* reporter described above was used to monitor T3SS gene expression. We first sought to test the level of T3SS expression via P*_vopN_-gfp* with titrated induction of ExsA. Addition of increasing concentrations of IPTG and theophylline resulted in increasing GFP production in the P*_tac-theo_-exsA* strain compared to the vector control, as expected (Fig. 3A). To test our hypothesis that LuxR regulates T3SS via another mechanism beyond regulation of *exsA*, we compared the Δ*exsA* Δ*exsB* strain to the Δ*exsA* Δ*exsB* Δ*luxR* strain. We observed that in the absence of LuxR there was a consistent increase in transcription from the *vopN* promoter compared to the presence of LuxR (Fig. 3A, compare yellow versus blue bars).

**Figure 3.**
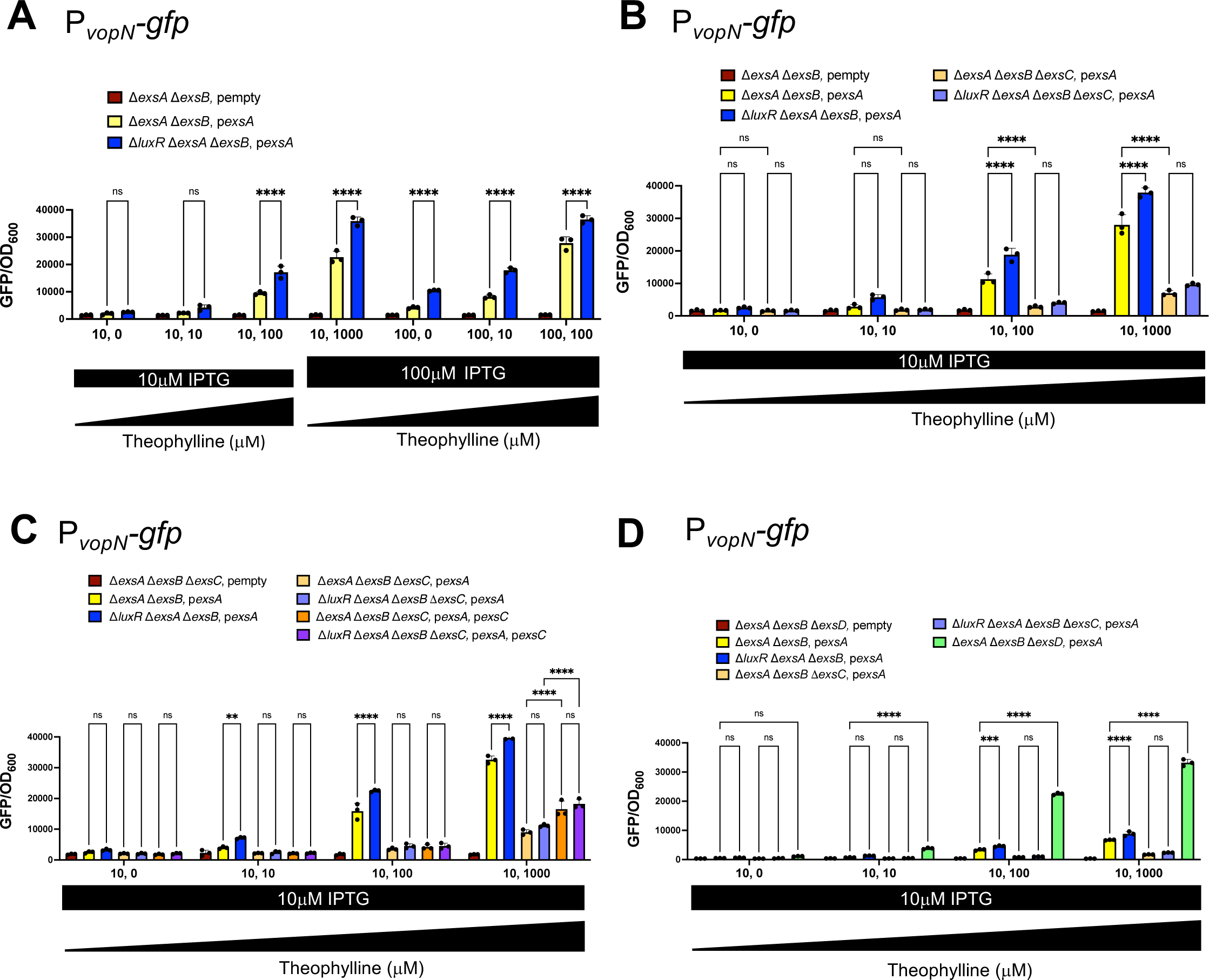
LuxR regulates T3SS genes through *exsB* and *exsC* loci. For all panels, genes *exsA* (plasmid pPP25: P*_tac-theo_-exsA;* denoted as p*exsA*) and *exsC* (plasmid pPP62: P*_tac-theo_-exsC;* denoted as p*exsC*) were induced via a P*_tac-theo_*-riboswitch^42, 43^ using either 10 μM or 100 μM IPTG in combination with 0, 10 μM, 100 μM, or 1000 μM theophylline. The empty vectors lack the *exsA* or *exsC* overexpression constructs. Error bars represent the standard deviation of means of *n*=3 biological replicates. Statistical significance between variables was determined using two-way ANOVA followed by Tukey’s multiple comparisons test on normally distributed data (Shapiro-Wilk test), *p <* 0.05.

To test our hypothesis that LuxR binds to the T3SS structural promoters I-IV to regulate transcription, we performed bioinformatic analysis of LuxR binding sites on the BB120 genome (43). However, no binding sites were predicted at the I, II, III, or IV promoters. On the other hand, our bioinformatic analysis identified multiple LuxR sites at the *exsB* and *exsC* promoters (Fig. 4A). To further test these results, we performed *in vitro* DNase I footprinting assay at the *exsC* and *exsB* promoters with LuxR. We observed protection at the two predicted sites closer to the *exsB* gene (Fig. 4A), and two overlapping predicted sites closer to the *exsC* gene as predicted in our bioinformatic analysis. Thus, we uncovered the *exsC* promoter as a putative second locus of LuxR-mediated regulation and repression of T3SS in *V. campbellii*. We constructed an *exsC* promoter fusion with a promoterless *gfp* cassette and assayed expression in the presence and absence of ExsA or LuxR. We observed that transcription from the *exsC* promoter was activated by ExsA and repressed by LuxR (Fig. 5A); complementation of either *exsA* or *luxR* restored expression to levels similar to the parent strain background. The pattern of regulation was further corroborated by RT-qPCR (Fig. 5B).

**Figure 4.**
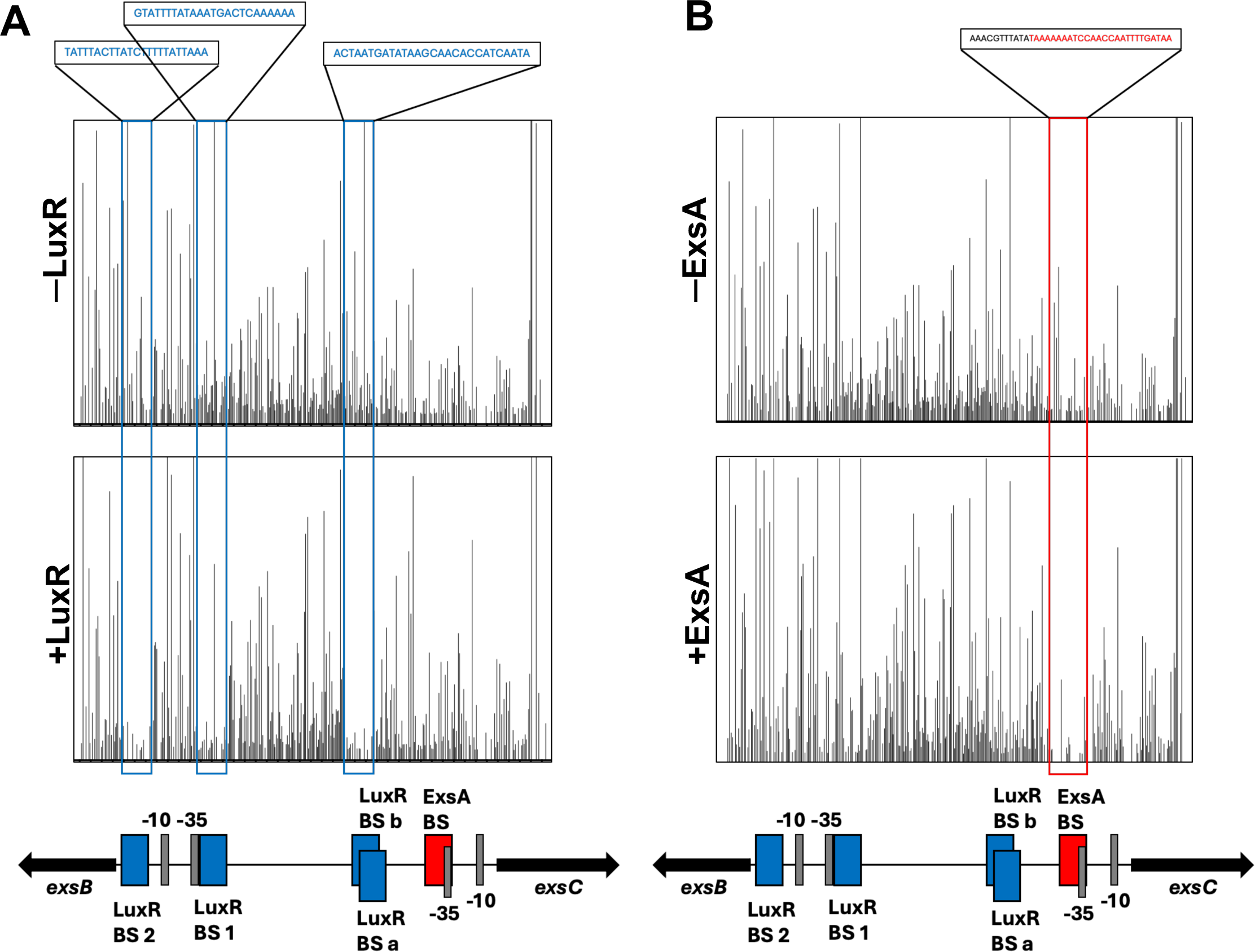
DNase I footprinting assays at the *exsC* and *exsBA* promoter region. **(A-B)** The probe is 425-bp long and labelled with HEX fluorophore dye at the 5’-end on the reverse strand and spans the *exsC* and *exsB* promoters. The final concentration of DNA probe was 20 nM. **(A)** DNase I-only treated probe (top) was compared with 0.25 μM LuxR- and DNase I-treated probes (bottom). **(B)** DNAse I-only treated probe (top) was compared with 1 μM ExsA- and DNase I-treated probes.

**Figure 5.**
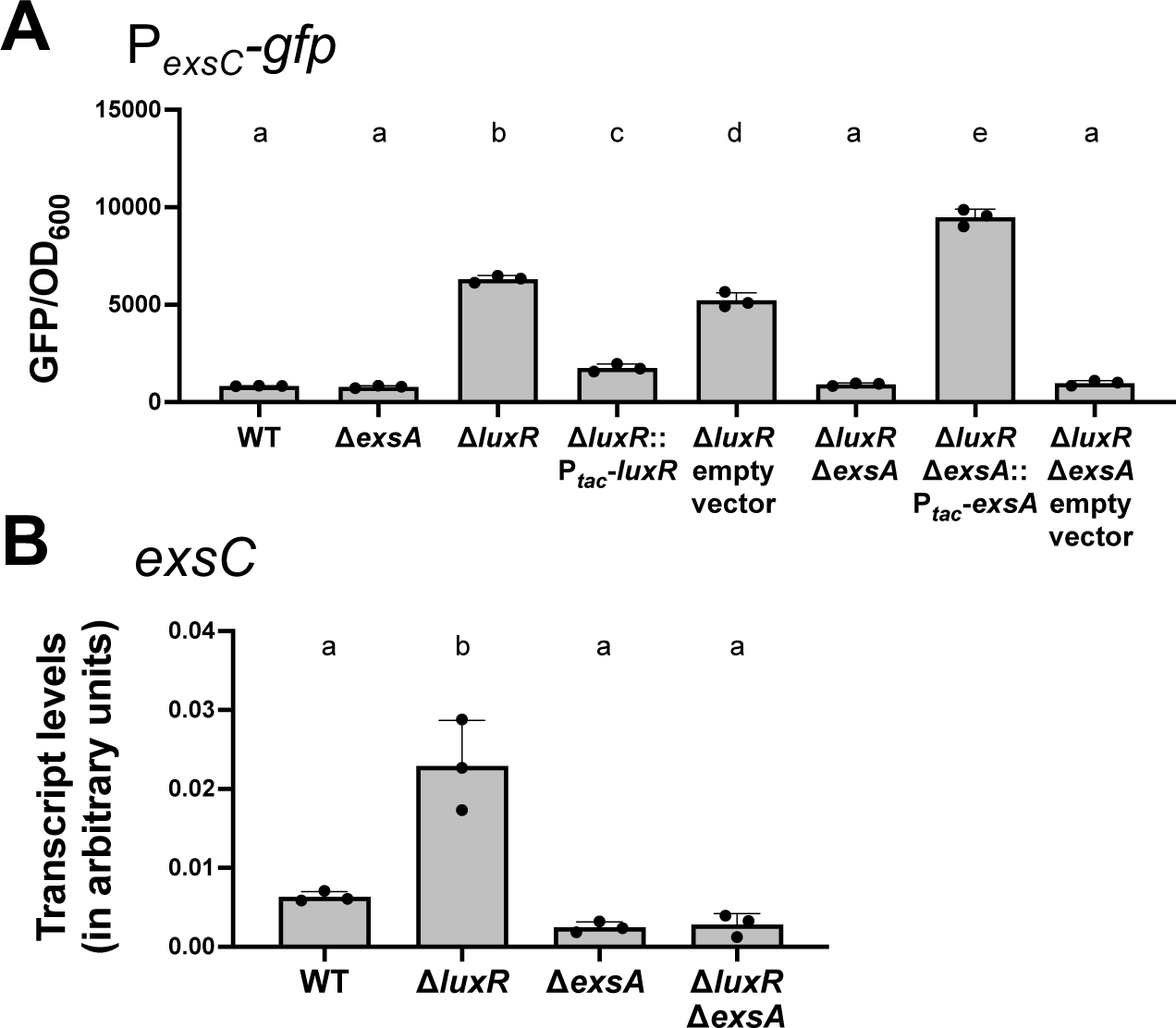
LuxR binds at the T3SS loci *exsB* and *exsC.* **(A)** GFP reporter assay with P*_exsC_-gfp* with isogenic strains containing plasmid pPP51. **(B)** RT-qPCR assay with isogenic BB120 strains to measure absolute transcript levels of *exsC* compared to internal control *hfq*. Error bars represent the standard deviation for triplicate biological replicates. A one-way analysis of variance (ANOVA) test was performed on normally distributed data (Shapiro-Wilk test) followed by Tukey’s multiple comparisons test. Different letters indicate significant differences between strains in pairwise comparisons (*p<*0.05; *n*=3).

To determine the role of the LuxR binding sites in the *exsBA* and *exsC* promoters, we constructed deletions and measured transcription using promoter-*gfp* fusions. Deletion of LuxR binding sites ‘1’ or ‘2’ at P*_exsB_* in a wild-type background did not de-repress *exsB* transcription, indicating that deletion of either site is not sufficient to eliminate LuxR repression of transcription from the *exsB* promoter (Fig. S4B). Further, deletion of both LuxR binding sites ‘a’ and ‘b’ (simultaneously) at P*_exsC_*was insufficient to alter *exsC* transcription (Fig. S4C). However, deletion of sites ‘1’ or ‘2’ in the P*_exsB_-gfp* reporter or sites ‘ab’ in the P*_exsC_-gfp* reporter (all in the Δ*luxR* background strain) led to significantly reduced GFP expression compared to intact promoters, suggesting that deletion of any of these sites impacts general transcription (Fig. S4B, S4C). It is possible that LuxR binding to all sites in the intergenic region between *exsC* and *exsBA* is required to exert repression of both promoters. From these results, we conclude that at HCD, LuxR binds to the *exsB* and *exsC* promoters but does not bind to the I-IV structural gene promoters under the conditions we have tested.

### LuxR represses ExsC, the anti-anti-activator of ExsA

In *P. aeruginosa* and *V. parahaemolyticus*, ExsC is a chaperone protein that binds the secretory protein ExsE during non-inducing conditions (Fig. 1A). During inducing conditions, ExsE is secreted through the T3SS machinery, which drives the formation of ExsC-ExsD complex. This event releases ExsA, which binds T3SS gene promoters and activates their transcription (25, 27). The activation of *exsC* expression by ExsA is a regulatory mechanism by which ExsA keeps its anti-activator ExsD bound and sequestered away from itself (25, 27). Thus, we hypothesized that LuxR repression of *exsC* would promote the ExsA-ExsD interaction and prevent transcription of the T3SS structural genes. To test our hypothesis, we first tested the effect of *exsC* deletion on P*_vopN_-gfp* expression. We observed that transcription from the *vopN* promoter was greatly diminished in the absence of ExsC (Fig. 3B, compare yellow versus light orange bars), which aligned with the predicted role of ExsC as an anti-anti-activator of ExsA. Next, to determine the role of LuxR in P*_vopN_-gfp* regulation in the absence of ExsC, we again utilized the P*_vopN_-gfp* reporter assay in combination with ectopic induction of *exsA* (P*_tac-theo_-exsA*) and compared GFP expression in a Δ*exsA* Δ*exsB* Δ*exsC* strain background in the presence and absence of *luxR*. We observed no significant difference in the presence or absence of LuxR (Fig. 3B, compare light orange versus light purple bars), indicating that the effects of LuxR regulation on *vopN* promoter expression were abrogated in the Δ*exsA* Δ*exsB* Δ*exsC* strain background. However, a significant difference in *vopN* transcription was observed between strains with *exsC* intact and *exsC* deleted (Fig. 3B, compare yellow and light orange bars). This indicated the importance of ExsC as a positive regulator of T3SS. As a complementation control for these results, we also tested the effect of overexpressing both ExsA and ExsC from the inducible P*_tac-theo_* promoter in the Δ*exsA* Δ*exsB* Δ*exsC* strain background (Fig. 3C, compare bright orange versus bright purple bars). We observed that induction of ectopic expression of ExsA and ExsC was able to partially complement activation of transcription from the *vopN* promoter (Fig. 3C, compare light orange versus bright orange bars, and light purple versus bright purple bars). However, even at the maximum levels of the inducers, the expression of P*_vopN_-gfp* did not reach the level in which *exsC* was intact. We hypothesized that this may be due to high levels of ExsD; ExsA also activates expression of *exsD*. Thus, even under conditions in which we overexpressed ExsA, we were also overexpressing its anti-activator ExsD, which may block full ExsA function. To test this hypothesis, we deleted the *exsD* gene in the Δ*exsA* Δ*exsB* strain, and once again utilized the P*_vopN_-gfp* reporter and ectopic induction of *exsA*. We observed that in the absence of *exsD*, transcription of P*_vopN_* was upregulated significantly more than the strains that had *exsD* intact (Fig. 3D, compare yellow versus green bars). From these results, we conclude that 1) LuxR represses the promoters of *exsB* and *exsC*, 2) ExsC activates T3SS gene expression, and 3) in the absence of *exsD*, T3SS gene expression is maximal.

### ExsA activates exsC but not exsBA

The ExsA binding site is highly conserved across Gram-negative species including in *Pseudomonas aeruginosa, Yersinia pestis*, *Yersinia enterocolitica*, *Photorhabdus luminescens*, and *Aeromonas hydrophila* (44, 45). Using the ExsA consensus binding motifs determined in earlier studies by DNase I footprinting assays in *Pseudomonas aeruginosa,* we performed an *in silico* search for ExsA binding sites across the T3SS pathogenicity island (Fig. S3B, S3C) (44, 45). We also performed differential RNA sequencing (dRNA-seq) to determine the precise position of transcription start sites (TSSs) in *V. campbellii* (Fig. S3D) and to precisely map ExsA binding sites relative to T3SS TSSs. We found predicted ExsA binding sites at the promoters of the four T3SS structural operons starting with *exsD*, *vopN*, *vscN,* and *vcrG* genes (Fig. S3C). Additionally, we found a predicted ExsA binding site at the *exsC* promoter; however, no predicted ExsA binding site was observed at the *exsB* promoter (Fig. S3C). To test our *in silico* results, we performed DNase I footprinting assay with 1 μM ExsA. The assay showed that ExsA indeed only binds at the *exsC* promoter at the consensus binding site spanning the −35 site and an A-rich region upstream of it (Fig. 4B). No binding was observed near the *exsB* promoter (Fig. 4B). Further, we tested the role of ExsA in the activation of transcription from the *exsB* promoter by assaying for GFP expression from a P*_exsB_*-*gfp* fusion reporter in Δ*luxR* and Δ*exsA* Δ*luxR* strains at HCD state. Even though deletion of *luxR* led to significantly increased GFP expression compared to the wild-type strain, no significant difference was observed in the case of Δ*exsA* Δ*luxR* strain compared to the Δ*luxR strain* (Fig. S4B). The assay showed that while LuxR repressed the *exsB* promoter, ExsA was not required for its activation. The absence of a predicted ExsA binding site at the *exsB* promoter was thus corroborated by our *gfp*-reporter assay.

We proceeded to test the impact of ExsA binding to its site in the *exsC* promoter. We constructed a P*_exsC_*-*gfp* fusion reporter with and without the ExsA binding site and assayed GFP expression in the Δ*luxR* strain background (Fig. S4A). Deletion of the ExsA binding site significantly reduced transcription of P*_exsC_*-*gfp* to levels similar to a wild-type strain and wild-type *exsC* promoter (Fig. S4C). From these data, we conclude that ExsA binding at the *exsC* promoter activates transcription of *exsC* through binding at a single site in the *exsC* promoter.

### LuxR alters ExsA binding at exsC

We sought to investigate the molecular mechanism by which LuxR regulates the *exsC* promoter. We initially tested binding patterns of LuxR and ExsA using *in vitro* gel shift assays with a probe corresponding to the *exsC* promoter. LuxR bound the *exsC* promoter substrate with shifts observed starting at 20 nM protein concentration (Fig. 6A). ExsA bound the *exsC* promoter with a higher affinity, with a shift in DNA migration observed starting at 7.81 nM protein concentration (Fig. 6B). The binding of ExsA at the *exsC* promoter shows multiple shifts observed at higher concentrations of the protein (Fig. 6B), possibly indicating DNA bending activity by ExsA. DNA binding by AraC family proteins as a dimer and subsequent bending of the DNA has been shown in previous studies of ExsA homologs in *P. aeruginosa* and *Y. pestis* (44). The DNA binding pattern of ExsA on the *exsC* promoter substrate suggests that ExsA binds the *exsC* promoter strongly. As negative controls, we performed gel shift assays with a probe corresponding to the *mutS* locus to test for LuxR and ExsA specificity. We observed no shift for either ExsA or LuxR at the control locus even at high concentrations of 3 μM and 500nM respectively (Fig. S5A, S5B). To further test for specificity of ExsA binding at the *exsC* promoter, we performed competitive gel shift assays wherein the fluorescently labelled *exsC* probe was competed with increasing concentrations of either unlabeled *mutS* probe or unlabeled *exsC* probe. We observed that ExsA binding at the labelled *exsC* probe (2 nM concentration) was not disrupted by the *mutS* probe even at a high concentration of 100 nM (Fig. S6A). On the other hand, competing the labelled *exsC* probe (2 nM concentration) with an increasing concertation of unlabeled *exsC* probe disrupted ExsA interaction at 20 nM and 100 nM concentrations (Fig. S6B). These results together show that binding of both ExsA and LuxR at the *exsC* promoter is highly specific.

**Figure 6.**
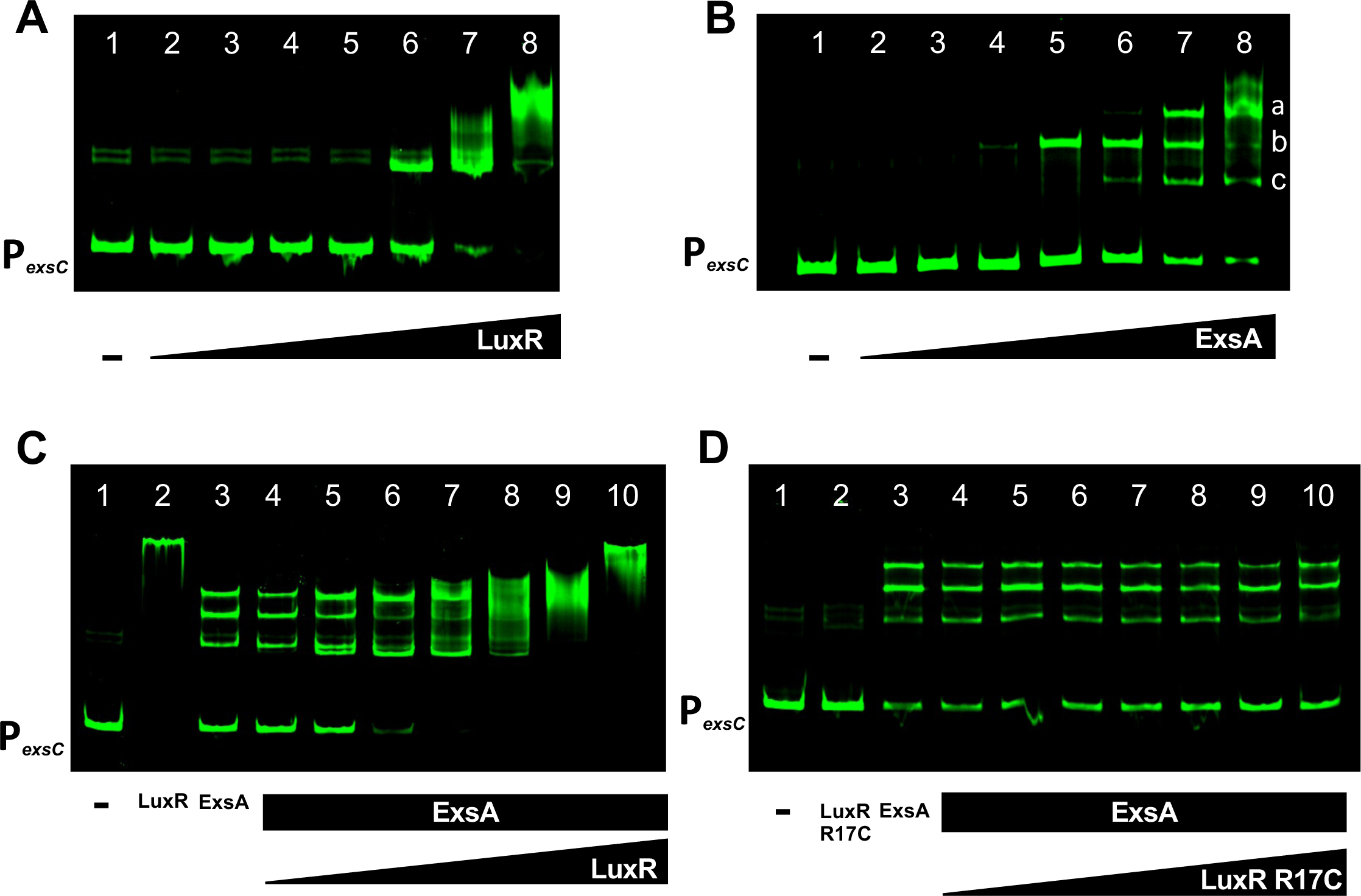
LuxR and ExsA bind the *exsC* promoter. **(A-D)** Electrophoretic mobility shift assays with a 200-bp DNA probe corresponding to the *exsC* promoter were assayed at a final concentration of 2 nM. Lane 1, DNA probe only control. **(A)** Lanes 2-8, 5-fold dilution series of LuxR from 500 nM to 0.032 nM. **(B)** Lanes 2-8, 4-fold dilution series of ExsA from 2 μM to 0.488 nM. Letters a, b, and c indicate different shifted band patterns. **(C, D)** Lane 2, LuxR or LuxR R17C alone at 700 nM and 1 μM concentrations, respectively. Lane 3, ExsA alone at 500 nM concentration. Lanes 4-10, ExsA is added at a constant concentration of 500 nM, followed by addition of either LuxR in a 2-fold dilution series from 700 nM to 10.94 nM, or LuxR R17C in a 2-fold dilution series from 1 μM to 15.625 nM.

Because we observed that LuxR and ExsA both bound at the *exsC* promoter, we hypothesized that LuxR represses *exsC* promoter by disrupting ExsA binding, which we previously observed to be required for *exsC* activation. To test our hypothesis, we performed competitive gel shift assays with LuxR and ExsA on the *exsC* promoter substate (Fig. 6C). LuxR disrupted the defined ExsA binding pattern starting 87.5 nM, despite ExsA being bound to the probe at 500 nM (Fig. 6C). We performed the same competitive gel shift assay but instead with DNA binding mutant LuxR R17C (43). No change in shift pattern was observed at increased concentrations of LuxR R17C (Fig. 6D). This result suggested that the disruption of the ExsA DNA binding patterns observed was due to LuxR-specific DNA interactions at the *exsC* promoter and not due to interactions between the proteins. We further performed competitive DNase I footprinting assays wherein we incubated a DNA probe spanning the *exsB* and *exsC* promoters with 1 μM ExsA followed by incubation with 0.25 μM LuxR (Fig. 7). We observed that ExsA remained bound at the *exsC* promoter even after addition of LuxR (Fig. 7). We have two hypotheses based on these results: 1) LuxR binding does not displace ExsA from its binding site in P*_exsC_* but disrupts DNA bending by ExsA. In this case, the active DNA conformation is not formed, and *exsC* transcription cannot be activated. 2) LuxR DNA binding does not affect ExsA DNA binding, and both proteins co-bind the promoter region, resulting in the observed gel shift and DNase I footprinting patterns. In this case, LuxR may block transcription by occluding another required transcription factor.

**Figure 7.**
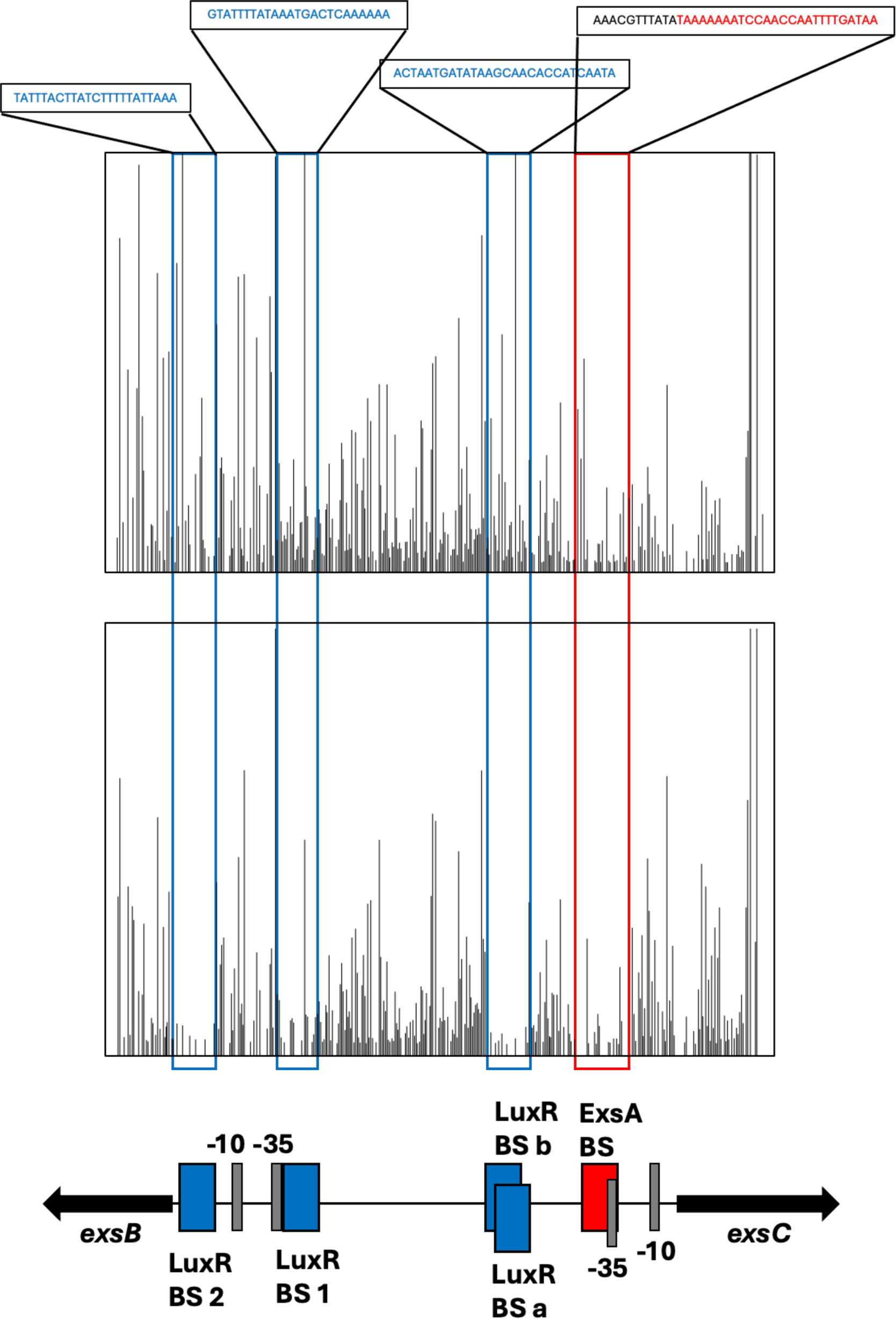
Competitive DNase I footprinting assay with LuxR and ExsA. The DNA probe is 425-bp long and labelled with HEX fluorophore dye at the 5’-end on the reverse strand and spans the *exsC* and *exsB* promoters. Final concentration of DNA probe used in reaction mix is 20 nM. DNAse I only treated probe (top) was compared with probe treated with 1 μM ExsA and 0.25 μM LuxR followed by DNase I treatment (bottom). Protection by ExsA was observed along the predicted ExsA binding site at the *exsC* promoter proximal to the −35 site. Protection by LuxR was observed along the predicted LuxR binding site at the *exsC* and *exsB* promoters.

## DISCUSSION

Our genetic epistasis and biochemical assays have uncovered a new regulatory role of the master QS transcriptional regulator LuxR in controlling T3SS expression: LuxR represses transcription of the anti-anti activator gene *exsC* (Fig. 8). Thus, our model proposes a dual mechanism of regulation controlling expression of the T3SS structural genes in response to changes in cell density (Fig. 8): at HCD, 1) LuxR represses transcription of *exsBA*, and thus the master transcription factor ExsA is not produced at HCD, and 2) LuxR represses transcription of *exsC*, and the decrease in ExsC protein promotes formation of the ExsD-ExsA complex and inactive ExsA protein. Our data suggest that this mechanism of regulation is a coherent type II feed forward loop wherein LuxR inhibits ExsA and ExsC separately, both of which act as activators of the T3SS structural genes (46). The regulatory pattern represents a distinction in the regulatory control of T3SS genes that may be divergent from that in other bacterial clades. There is apparent homology in sequence and function of *exs* genes in *Vibrio* and *Pseudomonas*, yet the synteny and regulatory patterns differ in numerous ways (Fig. S7) (25, 26, 33, 44). In both *V. parahaemolyticus* and *V. campbellii*, the expression of *exsB* and *exsA* are coupled and distinct from *exsCE*, whereas in *Pseudomonas*, *exsECB* are co-transcribed and distinct from *exsA* (18–20, 25, 33). In *V. parahaemolyticus*, ExsA activates its own expression from both the *exsB* and *exsA* promoters, yet in *V. campbellii*, ExsA does not activate its own expression based on the data we present in this study (33). This regulatory mechanism is more similar to *P. aeruginosa* in which transcription from P*_exsA_* is activated by global regulators Vfr, Fis, and VqsM but not ExsA (25). There are also similarities in regulatory patterns among the three organisms. ExsA activates the expression of its own anti-activator ExsD, thereby regulating its own activity through a feedback loop (25, 33). ExsC and ExsE are also activated by ExsA in *P. aeruginosa*, thus, it would be relevant to test whether *V. parahaemolyticus* ExsA also activates *exsCE*.

**Figure 8.**
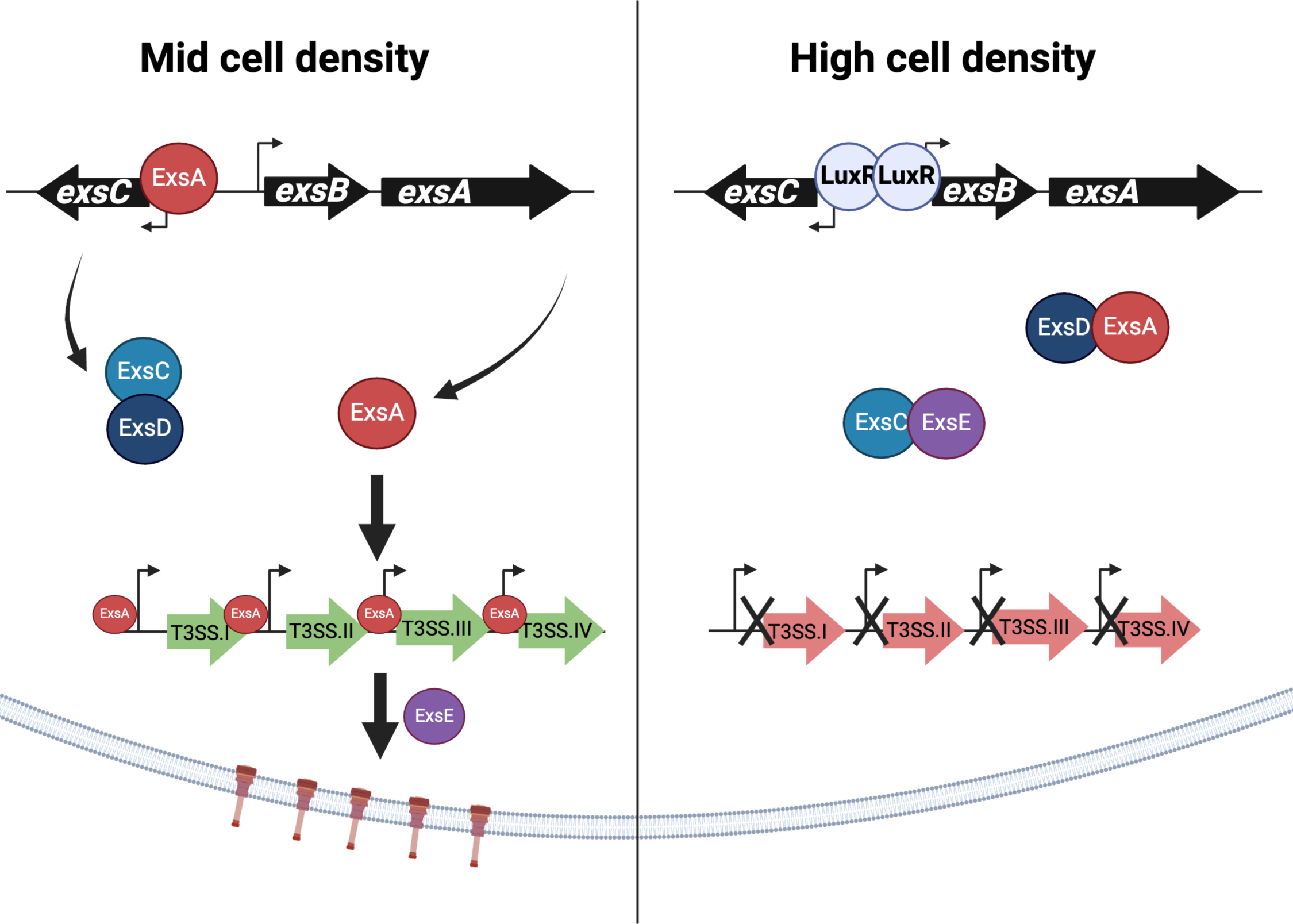
Proposed model of the dual mechanism of T3SS regulation by LuxR at mid cell and high cell densities in *V. campbellii*. At mid cell density, the absence of LuxR enables transcription of the *exsC* promoter by ExsA, which releases existing ExsA bound by ExsD. ExsA activates transcription of the four T3SS structural operons, enabling synthesis of the T3SS needles. At high cell density, LuxR represses transcription of *exsA* from the *exsBA* promoter and represses transcription of *exsC*. Lower levels of ExsC enable ExsD to bind to ExsA and prevent its activity as a transcriptional activator of T3SS genes.

During our examination of the divergent *exsBA* and *exsCE* promoters, we noted that the T3SS promoters differ from other σ70 promoters in that the spacer sequence between the −10 and −35 sites is longer than the canonical 17+/-1 bp length and tends to be closer to 20-22 bp in length instead (Fig. S3B, S3C). The *vopN* promoter has the exact −10 and −35 consensus sites for σ70 promoters, TATAAT and TTGACA respectively (Fig. S3B). The other T3SS promoters have approximately 50% homology to the consensus sites (Fig. S3B). The longer than canonical spacer region in the case of ExsA regulated promoters is reminiscent of MerR promoters. The MerR promoters have 19-bp spacer sequence between −10 and −35 sites (47). The binding of MerR to DNA under activating conditions causes a distortion in the DNA backbone so as to reorient the −35 and −10 sites that allows them to interact positively with RNA polymerase σ70 subunit (47). We hypothesize that ExsA binding is important to compensate for the weak −35 and −10 sites and the longer than usual spacer sequence. ExsA could be playing a similar role in which it brings the −35 site closer to the −10 site to match the consensus 17-bp spacer region and helps recruit RNA polymerase σ70 subunit to the ExsA dependent promoters. The *exsB* promoter with no predicted ExsA binding site differs from the other T3SS promoters in that it has the canonical 17+/-1bp spacer sequence between its −35 and −10 sites, also there is a conspicuous lack of the A-rich region upstream of the −35 site as is present in the other T3SS promoters with predicted ExsA binding sites.

The ExsA binding site has two important features, an A-rich region and the −35 site downstream of it with a 13bp gap in between (Fig. S3B, S3C). In *P. aeruginosa*, ExsA binds DNA as a monomer at a half site and recruits a second monomer to bind the other half site, thereby dimerizing post DNA binding (44). Separate binding events by ExsA monomers followed by dimerization leads to multiple shifts observed in gel shift assays with *P. aeruginosa* T3SS promoters including P*_exsC_*, P*_exsD_*, and P*_exoT_*(44). Thus, the multiple shifts we observed of ExsA binding at P*_exsC_* could be indicative of dimerization or polymerization of ExsA post-DNA binding, leading to DNA twisting or bending for transcription activation (Fig. 6B). The ExsA consensus binding site has been observed at all four structural promoters of the T3SS, indicating the expected necessary role of ExsA in activating transcription from these promoters (Fig. S3C).

We observed that deletion of individual LuxR binding sites at the *exsCBA* promoter region was insufficient to de-repress transcription from the P*_exsC_* and P*_exsB_* promoters (Fig. S4B, S4C). In a previous study, it was noted that global regulators LuxR and H-NS co-occupy and co-regulate transcription from 28 promoters responsible for the regulation of 63 genes including the *exsB* and *exsC* loci (31). At many of these promoters, multiple LuxR binding sites are present, and we have previously postulated that multiple LuxR binding sites function with redundancy to outcompete H-NS (31). We hypothesize that the incomplete de-repression of P*_exsC_* and P*_exsB_*promoters by deletion of only a single LuxR binding site could be due to the occupation of H-NS at the promoter regions and/or by the functional redundancy of multiple LuxR sites in the *exsBA exsC* intergenic region. Additionally, we observed in an earlier GFP reporter assay reporting on the transcription of *vopN* that Δ*luxR* Δ*hns* double mutant had significantly higher GFP expression compared to Δ*luxR* or Δ*hns* single deletion mutants (Fig. 2C). These results together could allude to the role of LuxR and H-NS in co-repressing the T3SS at HCD and specifically at P*_exsC_*.

The results of our studies here and combined with previous studies show that the QS system in *V. campbellii* shuts down the T3SS at HCD via two separate mechanisms to block ExsA activity: transcriptionally and post-translationally (Fig. 8). However, the evolutionary advantage for the repression of T3SS synthesis and activity at HCD is unclear. We speculate that expression of T3SS at low- to mid-cell density is advantageous at some stage during host infection. There is precedence for this; in *V. cholerae*, temporally controls virulence during infection (48). At late-stage infection, the *V. cholerae* QS system shuts down virulence genes and turns on genes that favor cell dispersal either to form new foci of infection in the intestine or to exit the host and re-enter the aquatic environment (48). We postulate that the *V. campbellii* QS system shuts down virulence in late-stage infection for similar reasons: dispersal of cells so as to exit the host and re-enter the aquatic environment in search of new host to colonize or to continue the planktonic lifestyle. Experimental assessment of these hypotheses may lead to a better understanding of the *V. campbellii* shrimp infection mechanisms and adaptation to different niche conditions.

## MATERIALS AND METHODS

### Bacterial strains and media

The *E. coli* S17-1λ*pir* strain was used for cloning purposes, and the *E. coli* BL21 (DE3) strain was used for overexpression and purification of all proteins. *Escherichia coli* strains were cultured at 37°C with shaking (250–275 RPM) in Lysogeny Broth (LB) medium with 100 mg*/*mL kanamycin, 100 mg*/*mL gentamycin, and*/*or 10 mg*/*mL chloramphenicol when selection was required. *V. harveyi* BB120 was recently reclassified as *Vibrio campbellii* BB120 (a.k.a., ATCC BAA-1116) (9). BB120 and derivatives were cultured at 30°C with shaking (250–275 RPM) in Luria Marine (LM) medium with 100 mg*/*mL kanamycin, 10 mg*/*mL chloramphenicol, and*/*or 50 mg*/*mL polymyxin B when selection was required. Plasmids were transformed into electrocompetent *E. coli* S17-1λ*pir* cells and subsequently conjugated into *V. campbellii* strains. *V. campbellii* exconjugants were selected using polymyxin B (50 U*/*mL). In case of strains transformed with inducible plasmid pPP25 and pPP62, 10μM IPTG, and 10μM, 100μM, and 100μM theophylline was used to induce the cells.

### GFP reporter assays

Endpoint GFP reporter assays were performed. Bacterial cultures were inoculated in 3 mL LM, 15mM MgSO4 and 5mM EGTA, and grown shaking at 30°C and 275 RPM overnight. 1mL of each culture was centrifuged for 3 minutes at 13000 rpm to pellet down cells. The cell pellets were resuspended in 1mL 1X saline PBS. Optical density was measured at OD_600_ nm wavelength using either the Biotek Cytation3 plate reader or the Synergy H1 plate reader. GFP signal was read at 485nm excitation wavelength and 528nm emission wavelength. Autogain function was performed for automatic adjustment of GFP signal per strain per plate assay. The GFP signal obtained was normalized to growth by dividing GFP read by OD_600_ read.

### Molecular and chemical methods

PCR was performed using Phusion HF polymerase (New England Biolabs) and Iproof HF polymerase (BioRad). All oligonucleotides were ordered from Integrated DNA Technologies (IDT). PCR products and plasmids were sequenced using Eurofins Genomics. Cloning procedures are available upon request. DNA samples were resolved using 1% agarose (1× TBE). Unless otherwise noted, data are plotted for triplicate independent experiments. Symbols on graphs represent the mean values, and error bars are standard deviations. Statistical analyses were performed with GraphPad Prism version 10.2.0. Additional information about statistical analyses is included in the figure legends. CRISPRi knockdown constructs were generated as described (41) with targeting small guide RNAs against *exsC*, *exsD*, and *exsE*.

### Construction of deletion/epitope-tagged strains

All *V. campbellii* BB120 and KM669 derivative strains in this study were constructed following a previously published technique (49). Briefly, the pRE112 suicide vector was used to construct unmarked deletions or insertions of epitopes in which 1000 bp of upstream and downstream flanking sequence was cloned into pRE112. The pRE112 derivatives were conjugated into *V. campbellii* and selected on chloramphenicol to induce chromosomal recombination of the plasmid. Subsequently, the plasmid was excised via counterselection on 15% sucrose. Cells in which the plasmid excision yielded a non-WT locus were detected via colony PCR. All gene deletions were confirmed by DNA sequencing through Eurofins.

### Expression and purification of LuxR protein

LuxR, His-LuxR and FLAG-LuxR were purified as previously described (35, 43, 50) in BL21(DE3) cells containing either pJV079 (wild-type *luxR*) or pJV206 (*luxR* R17C).

### Expression and purification of ExsA protein

ExsA was purified by overexpressing N terminal 6X His tagged ExsA (pPP43 vector ordered from BioTwist) in *E. coli* BL21(DE3) cells. The purification protocol is a modified version of a previously described protocol (28). The cells were initially grown at 37°C to an OD_600_ = 0.6–0.8 and ExsA overexpression was induced by using 1 mM IPTG for 4 h. Induced cell pellets were resuspended in lysis buffer (20 mM Tris–HCl pH 8.0, 500 mM NaCl, 20 mM imidazole, 1×protease inhibitors, 1 mM PMSF, 0.2 mg/ml DNaseI (GoldBio), 1× FastBreak (Millipore)) and incubated at room temperature for 35 min. Tween 20 was added to a final concentration of 0.5% following lysis. The protease inhibitor mix included the following: 0.07 mg/ml phosphoramidon (Santa Cruz), 1.67 mg/ml AEBSF (DOT Scientific), 0.07 mg/ml pepstatin A (DOT Scientific), 0.07 mg/ml E-64 (Gold Bio), and 0.06 mg/ml bestatin (MPbiomedicals/Fisher). Clarified lysate was loaded onto a 5 mL His-Trap Ni-NTA column (GE Healthcare Life Sciences) using an Akta Pure (GE Healthcare Life Sciences). Protein was eluted from the column using a linear gradient of elution buffer (25 mM Tris–HCl pH 8.0, 500 mM NaCl, 500 mM imidazole). Fractions were analyzed by SDSPAGE to confirm the presence of 6xHis-ExsA and pooled together. Pooled fractions were concentrated using 10 kDa cutoff centrifugal filters (Sartorius) and dialyzed overnight in storage buffer (20mMTris–HCl pH8.0, 500mM NaCl, 1 mM DTT, 0.5% Tween 20). Dialyzed protein was aliquoted, snap frozen in liquid N2, and stored at −80◦C.

### Electrophoretic mobility shift assays

Primers to amplify over 200bp of the promoter regions of exsB and exsC were ordered from IDT where in the 5’ end of one primer among the pair was labelled with IRD800CWN. PCR amplification was used to produce promoter substrates. Labelled and amplified DNA probe was incubated for 30 min in a 15mL reaction mixture containing either 20X stock ExsA binding buffer (20 mM Tris [pH 7.5], 100 mM KCl, 2mMdithiothreitol [DTT], 2 mM EDTA, 5% glycerol), or 10X stock LuxR binding buffer (10 mM HEPES [pH 7.5], 100 mM KCl, 2 mM dithiothreitol (DTT), 200 mM EDTA), 10 ng/mL poly(dI-dC), 0.1mg/ml bovine serum albumin (BSA), and the desired protein diluted in either ExsA dilution buffer (1X stock ExsA binding buffer), or LuxR dilution buffer (20 mM Imidazole [pH 7.5], 3000 mM NaCl, 0.5 mM EDTA, 1 mM DTT, and 5% glycerol). The reaction mixtures were separated on 6% TGE (25 mM Tris, 0.25 M glycine, 1 mM EDTA)-polyacrylamide native gels. Binding reactions were performed using either 20 nM labelled DNA or 20 nM unlabeled bringing the final concentration of DNA probe to 2nM in the reaction. Poly(dI-dC) (10 ng/mL, Sigma) was used as non-specific competitor DNA in all cases. For competitive binding experiments, reactions were supplemented with the secondary protein in binding buffer and incubated at room temperature for an additional 30 min.

### qRT-PCR, RNA extraction, and differential RNA-seq

Strains were inoculated in 5 mL LM and grown overnight shaking at 30°C at 275 RPM. Each strain was back-diluted 1:1000 in LM and 15mM MgSO_4_ and grown shaking at 30°C at 275 RPM until they reached an OD_600_ = 1.0. 2mL cells were collected by centrifugation at 3700 RPM at 4°C for 10 min, the supernatant was removed, and the cell pellets were flash frozen in liquid N_2_ and stored at −80°C. RNA was isolated from pellets using a TRIzol/chloroform extraction protocol and treated with DNase via the DNA-free™ DNA Removal Kit (Invitrogen) as previously described (51). Quantitative reverse transcriptase real-time PCR (RT-qPCR) was used to quantify transcript levels of T3SS genes in different regulatory conditions and was performed using the SensiFast SYBR Hi-ROX One-Step Kit (Bioline) according to the manufacturer’s guidelines. Primers were designed to have the following parameters: amplicon size of 100 bp, primer size of 20–28 bases, and melting temperature of 55–60°C. All reactions were performed using a Step One Real PCR system with 0.4 μM of each primer and 200 ng of template RNA (20 μl total volume). All RT-qPCR experiments were normalized to the internal standard *hfq* gene. The ΔΔ*C_T_* or standard curve methods were used to analyze data from at least three independent biological replicates with two technical replicates each.

For dRNA-seq, cells were collected as described above but without MgSO_4_. The cDNA libraries were constructed as described previously (52) by Vertis Biotechnology AG (Freising, Germany) and sequenced using an Illumina NextSeq 500 machine in single-read mode (75 bp read length). Reads were mapped and transcription start sites determined as described previously (52). The raw, demultiplexed reads and coverage files have been deposited in the National Center for Biotechnology Information Gene Expression Omnibus with accession code GSE182898.

### DNase I footprinting assay

The promoter region for transcription activation of the *exsC* and *exsB* genes with LuxR and ExsA binding sites in *V. campbellii* BB120 (425 bp) was PCR-amplified using a forward primer with a 5′-FAM fluorescent tag (IDT) and a reverse primer with a 5′-HEX fluorescent tag (IDT). DNase I (NEB) concentration was optimized by assessing the activity of DNase I across a range of concentrations (1x-1024x, diluted with water), and the optimal concentration for the DNase I enzyme and buffer conditions was determined to be 128-fold dilution. Footprinting reactions were set up in the following conditions: 50 mM Tris-HCl pH 7.5, 10 mM MgCl_2_, 50mM KCl, 0.1 mg/mL BSA, 1mM DTT, 5% glycerol, 20 nM DNA probe, and either no protein or 1 µM ExsA or 0.25μM LuxR or both. Reactions were incubated at room temperature for 15 minutes. Subsequently, 5μL of DNase I (diluted 128-fold) was added to each reaction and incubated for 15 minutes at room temperature. To stop the digestion reaction, 25 µL 0.5 M EDTA pH 8.0 was added. DNA fragments were recovered using Qiagen MinElute PCR clean up columns and eluted into black Eppendorf tubes. DNA fragments were analyzed with Genewiz Azenta Life Sciences using their Fragment analysis service and LIZ 500 DNA standard ladder. Peak Scanner software v1.0 was used for data analysis.

### Generation of ExsA consensus binding site

Multiple sequence alignment for the ExsA binding site sequences was extracted from the previously published ExsA-dependent promoter sequences and putative T3SS promoter sequences from *Pseudomonas aeruginosa*, *Photorhabdus luminescens* and *Aeromonas hydrophila* (28) using which a position weight matrix (PWM) was constructed. The PWM was searched against the *Vibrio campbellii* strain BB120 genome sequence using the FIMO (FInd MOtif) tool (53) from MEME suite version 5.1.0. Using the search hits with qval ≤0.01, the process of PWM construction followed by FIMO search was repeated iteratively until no more additional hits could be found. The sequence conservation among the significant matches to the ExsA motif was visualized as a sequence logo generated using Seqlogo version 2.8 (54).

## Acknowledgments

The authors thank Dr. Timothy Yahr, Dr. Daniel Kearns, and the van Kessel lab for comments on the manuscript and helpful discussions. We also thank Chelsea Simpson and Victoria Lydick for excellent technical support. Research reported in this publication was supported by: 1) the National Institute of General Medical Sciences (NIGMS) of the National Institutes of Health (NIH) under award numbers R35GM124698 to JVK, The content is solely the responsibility of the authors and does not necessarily represent the official views of the National Institutes of Health; 2) the National Science Foundation Graduate Research Fellowship Program to LJG. K.P. acknowledges support from the DFG (SPP2389, project-ID 503931087 and EXC 2051, project-ID 390713860) and the European Research Council (ArtRNA, CoG-101088027).

